# Thermally driven sex reversal reveals divergent sex determination dynamics in wild viviparous reptile populations

**DOI:** 10.64898/2026.06.25.734438

**Authors:** Carles Ferre-Ortega, Paul A. Saunders, Shane A. Richards, Christopher Burridge, Luisa J. Fitzpatrick, Peta Hill, George D. Cunningham, Geoffrey M. While, Tariq Ezaz, Erik Wapstra

## Abstract

Climate change can threaten population viability by disrupting sex ratios in species whose sex is influenced by temperature. While species with sex chromosomes were historically considered immune, in some species, temperatures can override genetic sex determination via sex reversal, leaving them vulnerable to climate-driven sex ratio shifts. The Tasmanian spotted snow skink (*Carinacincus ocellatus*), a viviparous reptile with an XX/XY system, provides a compelling case study. While laboratory studies demonstrated that extreme thermal conditions induce female-to-male sex reversal (XX males), its occurrence in the wild remains unexplored, limiting our understanding of actual climate impacts. Integrating 23 years of phenotypic and genetic sexing data across two climatically distinct populations, we provide the first evidence of sex reversal in a wild viviparous reptile. XX reversal occurred in both populations, affecting up to 23.5% of XX births in the warmer population, and was associated with colder minimum daily temperatures. Despite high birth rates in some years, sex-reversed adults were rare. We also identified putative XY females, suggesting bidirectional sex reversal and reinforcing the extreme plasticity of reptilian sex determination. Ultimately, sex reversal could act as an evolutionary trap, potentially compromising population viability as climate instability increases.

## 1. Introduction

In species with separate sexes, males and females are typically produced in approximately equal proportions due to negative frequency-dependent selection [1]. However, deviations from parity can occur when producing offspring of a particular sex enhances parental fitness, as predicted by sex allocation theory [2]. Many organisms achieve such optima through environmental sensitivity (e.g., Temperature-dependent Sex Determination, TSD) [3]. Rapid environmental shifts, however, can decouple these cues from the benefits they were selected to generate, threatening population viability before compensatory mechanisms have time to evolve. Skewed sex ratios can intensify same-sex competition, elevating aggression levels and altering mating systems [4–6]. Moreover, male-biased populations may experience reduced reproductive output due to scarcity of females [7], whereas female-biased populations may exhibit enhanced reproductive rates in polygynous systems, although extreme female-skews can eventually precipitate population collapse [8,9]. Persistent imbalances can also promote inbreeding, erode genetic diversity, and reduce adaptive potential, thereby elevating the risk of population decline or extinction [8,9].

Anthropogenic climate change has intensified the risk of biased sex ratios in species with known thermal sensitivity [10–12]. In TSD species, sex ratio skews have already been associated with warming climates [12–14]. Species in which sex determination is governed by sex chromosomes (Genetic Sex Determination, GSD) were historically considered exempt from these thermal effects [15]. However, GSD and TSD are now considered ends of a continuum of sex determining modes rather than being mutually exclusive or dichotomous [16,17]. In some species, extreme temperatures can override the genetic influence on sex determination through sex reversal — a process occurring when the production of sex-determining factors is temperature-sensitive (Figure 1) [11,16,17].

**Figure 1:**
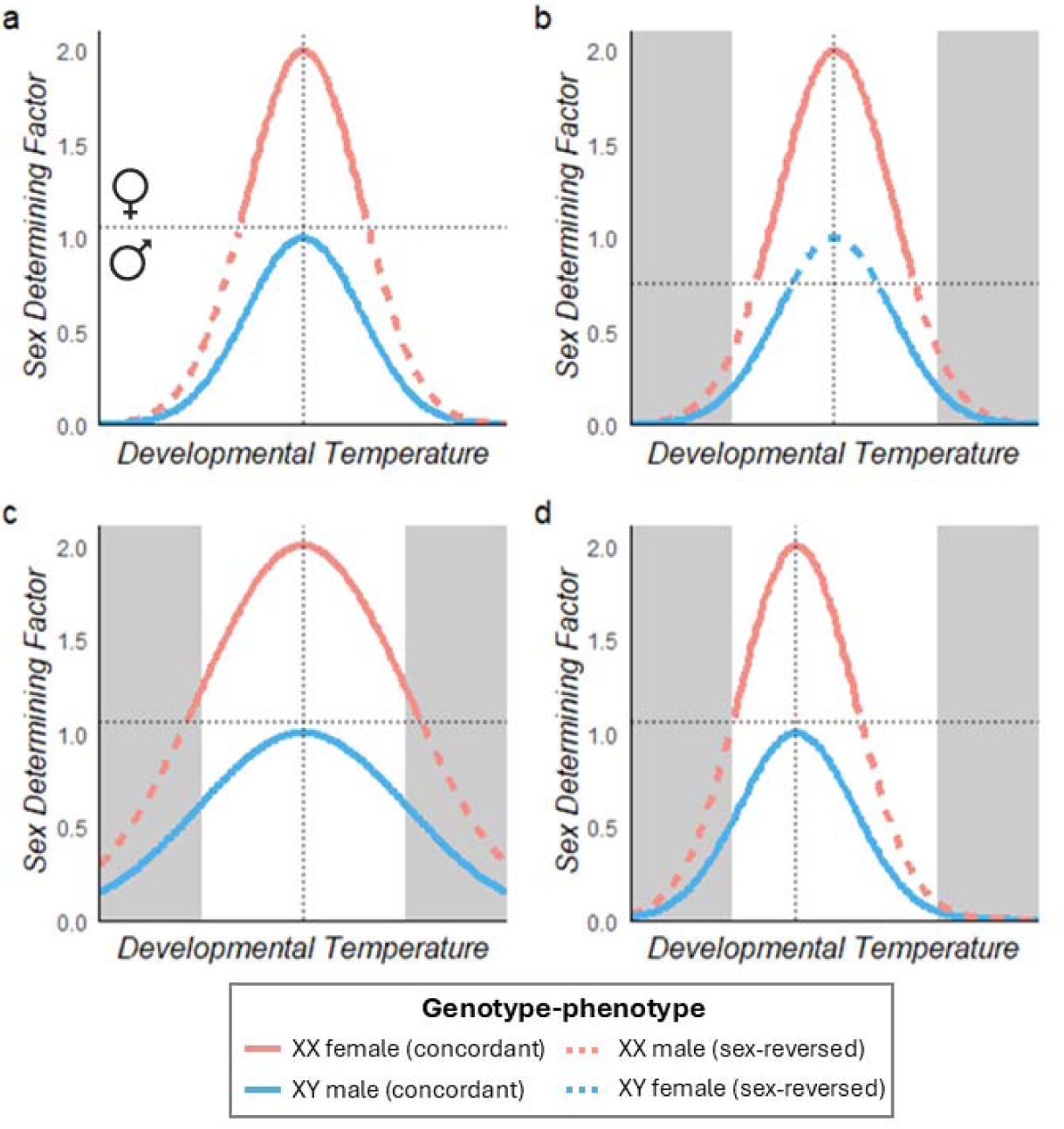
Mechanistic threshold model for sex determination, adapted from Quinn et al. [16]. (a) Sex is determined by the expression of a temperature-sensitive sex-determining factor (SDF). An individual’s expression curve depends on the temperature of peak SDF expression (vertical dotted line), the breadth of its reaction norm across developmental temperatures (width of the curve) and the developmental threshold for sex determination (horizontal dotted line). In this example where SDF is X-linked, XX individuals (red lines) naturally exhibit twice the baseline SDF of XY individuals (blue lines), typically exceeding the threshold for female development (continuous red lines). However, extreme developmental temperatures inhibit SDF expression, dropping XX individuals’ expression below the threshold and producing sex-reversed males (red dashed lines). Within the same range of developmental temperatures (unshaded areas), individual or population-level sex expression can vary due to: (b) changes in the sex determination threshold, altering the probability of XX males and allowing XY females to occur (blue dashed lines); (c) modifications in the width of the reaction norm; and/or (d) changes in the peak expression temperature. (*Equivalent dynamics apply to ZZ/ZW or dominant Y/W systems*).

The fitness consequences of sex reversal vary widely across taxa. While sex reversal has been associated with poor condition and physiological stress in juvenile agile frogs (*Rana dalmatina*) [18,19], it appears neutral in Nile tilapia (*Oreochromis niloticus*) [20] or even beneficial in the central bearded dragon (*Pogona vitticeps*), where sex-reversed females exhibit enhanced fecundity [21,22]. These fitness outcomes dictate the evolutionary trajectory of the sex-determining system. If sex-reversed individuals gain a fitness advantage, they can drive the transition from GSD to TSD; for instance, matings between XX males and XX females produce exclusively XX offspring, potentially leading to the loss of the Y chromosome and fixing the population as a TSD system [21,23]. Conversely, if sex reversal compromises individual fitness, it risks becoming an evolutionary trap, producing maladaptive phenotypes that may drive demographic collapse. Understanding how sex reversals shape population trajectories requires examining their occurrence and impacts on individual fitness through ontogeny across a broader range of species.

While sex reversal has been documented across a wide range of vertebrate taxa [24–26], reptiles have emerged as particularly valuable study systems. This matches their importance for the study of sex determination and sex chromosome evolution – possessing TSD, GSD, and both heteromorphic and homomorphic extremes of XY and ZW sex chromosome systems across related taxa [27,28]. Research on two oviparous species — the eastern three-lined skink (*Acritoscincus duperreyi*) and *P. vitticeps* — has provided foundational insights into the mechanisms underlying temperature-induced sex reversal [21,29].

Specifically, in *A. duperreyi*, reversal rates correlate with altitudinal thermal gradients [30], whereas in *P. vitticeps*, they are related to local genetic background [31]. Notably, these studies also provided the first evidence of positive fitness effects in sex-reversed individuals [21,22,32], with implications for the selective maintenance of sex reversal. However, we only have insights from these two oviparous species at present, despite the diversity in reptilian sex chromosome and life histories, which may influence our understanding of the likelihood and consequences of sex reversal.

The Tasmanian endemic spotted snow skink (*Carinascincus ocellatus*) is a viviparous reptile with XX/XY sex chromosome system [33,34] that offers promising opportunities to advance our understanding of sex determination and sex reversal. Long-term field and laboratory studies on a relatively warm lowland population have shown that offspring phenotypic sex ratios are temperature sensitive — strongly male-biased when gestational temperatures are colder, and to a lesser extent, female-biased when they are warmer [35–38]. Although previously interpreted as TSD, sex reversal has recently emerged as a plausible mechanism underlying these shifts, following documentation of sex chromosomes [33,34] and female-to-male sex reversal (i.e., XX males) under sustained lower developmental temperatures or restricted maternal basking opportunities in the laboratory [39]. Unexpectedly, a similar observation was made in the laboratory for females from a colder highland population despite offspring sex ratios maintaining parity across temperatures at that natural location [36–39]. This suggests that a single sex-determining system (GSD with thermal sensitivity) operates across the entire species, but its expression in the wild may be modulated by a combination of local adaptation and environmental factors, highlighting its ecological and evolutionary complexity.

To resolve sex reversal complexity and determine whether it actively shapes wild populations of *C*. *ocellatus*, we investigated its natural occurrence and frequency using a long-term dataset from two climatically distinct highland and lowland populations. This dataset encompasses over two decades of phenotypic sex records from wild offspring, coupled with targeted genetic sexing collected over several years. Using this dataset, our investigation addresses four aims. First, we quantify the natural occurrence of sex reversal in offspring cohorts across these climatic extremes. Second, we identify the environmental drivers of this phenomenon by testing the association between sex reversal and temperature, incorporating both maximum and minimum daily temperatures to comprehensively assess thermal effects. Third, we model 23 years of annual phenotypic sex ratios against these same thermal variables to determine whether temperature-induced sex reversal is concordant with population-level sex ratio shifts. This allows us to assess whether the thermal signature of sex reversal aligns with observed phenotypic biases, distinguishing it from shifts in the underlying genetic sex ratio (e.g., due to sex-biased mortality). Finally, we screen adult cohorts across eight populations to determine whether sex-reversed offspring persist to maturity and potentially contribute to the breeding population.

## 2. Material and methods

### Field and husbandry protocols of offspring analysis

We utilised two field populations of *C. ocellatus* at the extremes of the species’ climatic range which have been studied for more than two decades [36–38,40–42]. The first population, the ‘warm’ lowland population, was located at Orford (42°34’ S, 147°52’ E; 30 m a.s.l.), and the second population, the colder highland population, was located at the Central Plateau (41°51’ S, 146°34’ E; 1,150 m a.s.l.; Figure S1). In both, individuals reach maturity after 2–4 years and produce litters of one to eight offspring, with some variability between populations [38,43,44]. Ovulation is synchronized among females and consistent between years [36,38]. Gestation is 3–4 months long and is dependent on maternal basking opportunities [38,41,42,45].

Each season, the majority of gravid female skinks (∼90-95%, estimated based on field observations and long-term knowledge of population structures and densities) were systematically captured at each population at the end of their gestation period. This occurred in late December in the lowland population, as site temperatures were warmer and the gestation season started earlier [41], and in mid to late January in the highland population, well after the period when sex determination was sensitive to temperature [37,38,46].

Following capture, individuals were transported to University of Tasmania’s laboratories and maintained under standardised conditions until they give birth. Specifically, females were housed in individual terraria (30 x 20 x 15 cm) in a temperature (∼16 DC) and light (14h L: 10h D) controlled room. During the hours of 8 am to 6 pm, lizards had access to basking lights that provided a thermal gradient of 16–40 DC, allowing thermoregulation to preferred body temperatures [40,47]. At birth, each offspring was phenotypically sexed (see below) and toe-clipped for permanent identification. If not performed at birth, captured adult females were toe-clipped to facilitate identification during subsequent field recaptures, as they can live more than 10 years [44,48]. In most years, a small portion of the juvenile and adult tails were collected and preserved in ethanol for later DNA analysis. Following birth, all adult females were returned to their point of capture with offspring released randomly at specific designated sites within the broader source population.

### Temperature data collection

Temperature data for each site were obtained from two weather stations within the Australian Bureau of Meteorology network that were closest to our lizard populations. The Liawenee station (station number 96033; 41.90° S, 146.67° E, 1057 m a.s.l.) was used to extract highland population temperatures, while lowland population temperatures were extracted from the Aubin Court station (station number 92027; 42.55° S, 147.88° E, 14 m a.s.l.). Following previous analyses on sex ratio and sex determination in this species, the analysis of temperature data focussed on the 6-week period during which embryos’ sex was temperature sensitive [35,37,46]. This period spanned from October 1st to November 15th for the lowland population and from October 15th to November 30th for the highland population.

### Offspring phenotypic and genotypic sexing

Assessing evidence of biased sex ratios and sex reversal in the field required integration of three sources of information: (i) measurements of offspring phenotypes, (ii) an assessment of reliability of phenotypic sexing, and (iii) offspring genotypic sex.

Phenotypic sex in *C. ocellatus* was assessed through hemipenis eversion, which is a well-established method in reptiles [21,49] and *C. ocellatus* specifically [35,36,39,40]. Phenotypic sexing of neonates was performed within a few days of birth by the same assessor (EW) across all 23 years under identical conditions. While phenotypic sex classification is highly repeatable, misclassifications can occur. We incorporated empirically derived misclassification rates from Hill et al. [39] who exposed 100 gravid females from lowland and highland populations to varied thermal regimes. The same assessor (EW) phenotypically sexed the 418 resulting offspring using a blind repeat-assessment protocol. Discrepancies were rare: the probability of falsely scoring a phenotypic male as female was estimated at just 0.2%, and scoring a phenotypic female as a male was 2.3% [39]. These probabilities were integrated into our statistical models to ensure the reliability of our wild sex reversal estimates (see below).

We genetically sexed offspring of *C. ocellatus* (phenotypic males and females) to identify individuals with a mismatch between their genotypic and phenotypic sex (i.e., sex reversed). DNA was extracted from tail tips (<5 mm) by incubating them in 200 µL of 5% Chelex-100 solution with 40 µg of Proteinase K overnight at 56 °C, after which the supernatant was stored at -20 °C. Determination of genetic sex was conducted using a hierarchical PCR screening approach based on three markers developed by Saunders et al. [50] to distinguish *C. ocellatus* with XY and XX genotypes. In the primary screen, all samples were genotyped using Primer Set 1; individuals displaying a genotype-phenotype mismatch were then re-tested with Primer Sets 2 and 3 to ensure reliability. All individuals identified as sex reversed demonstrated a consistent mismatch across all three primer sets.

We genetically sexed offspring from the 2014, 2015, 2018, 2019, and 2022 cohorts for both populations. To ensure robust statistical power and adequately captured sex reversal dynamics given the higher sex ratio variation and smaller annual sample sizes from the lowland population (annual mean: lowland = 196.9, highland = 308.0), we additionally included the lowland cohorts from 2020 and 2021. These specific years were selected to capture some of the most pronounced male- and female-skewed phenotypic sex ratios observed in the dataset (Figure 2a). Samples collected before 2014, as well as those from 2016 and 2017, were excluded from the analysis due to poor DNA quality. On average, sampling completeness exceeded 95% of offspring per year. The lowest coverage occurred in the 2015 lowland cohort (78 %), where the loss of several DNA samples prior to screening limited the genotyped cohort to 61 of 68 phenotypic males and 38 of 56 phenotypic females.

**Figure 2:**
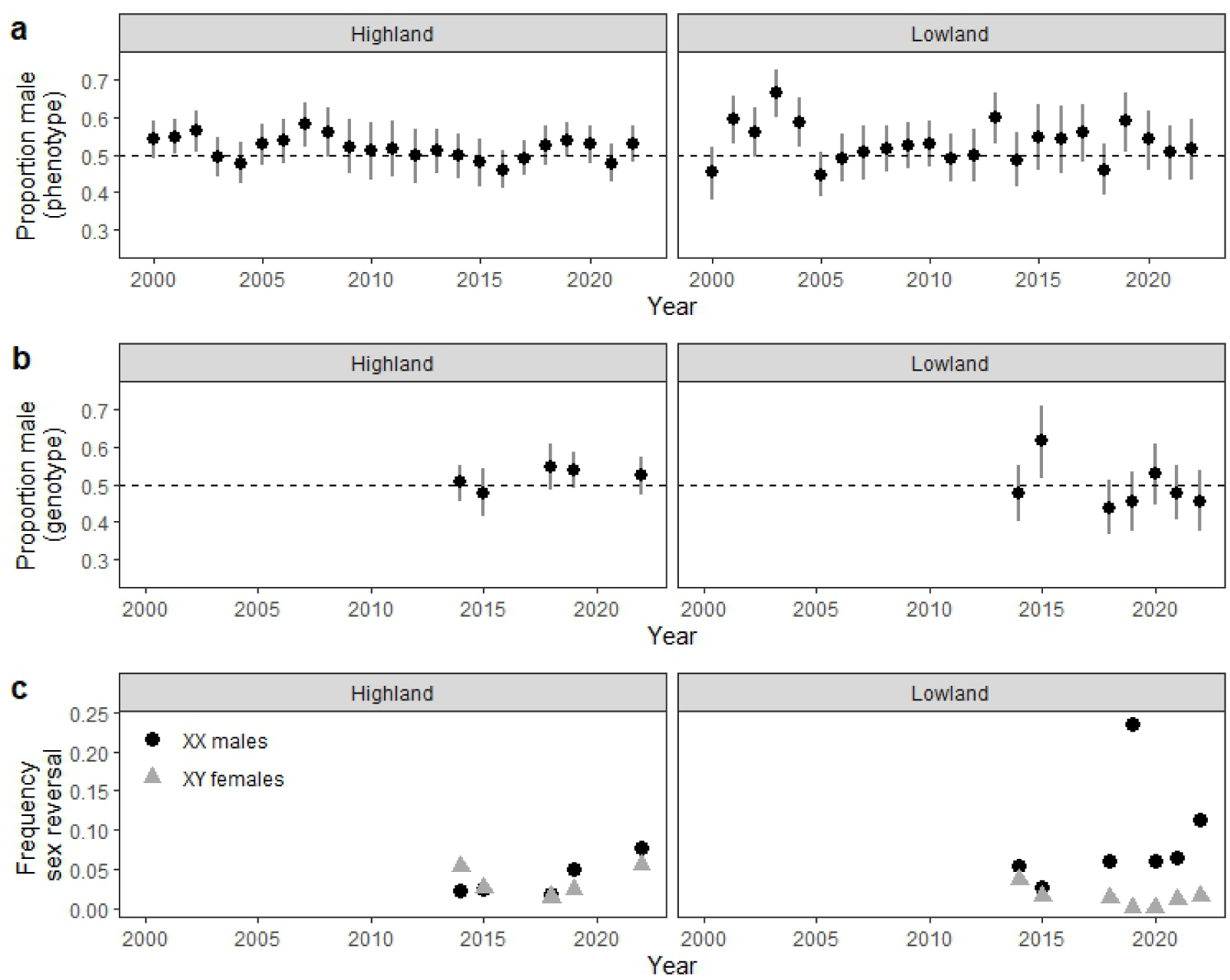
Annual sex ratios and sex reversal frequencies. Mean proportion per clutch of a) phenotypic males and b) genotypic males (XY) for highland and lowland populations. Bars indicate 95% confidence intervals; dashed lines denote a 50:50 ratio. c) Proportion of sex reversed individuals relative to the annual count of their respective genotypic sex (XX males = circles; XY females = triangles). *Note: The 2015 lowland male skew in (b) is a sampling artifact due to disproportionate pre-screening loss of phenotypic females*.

### Modelling temperature effect on genotypes and cases of sex reversal

Annual variation in the phenotypic sex ratio could at least be in part a consequence of sex reversal or annual variation in the genotypic sex ratio (e.g., due to sex-bias heat tolerance) [10]. We used a binomial regression approach to estimate both the probability an offspring was a genetic male (g) and the probability it exhibited sex reversal according to its genetic sex (r_g_). This framework allowed the inference of sex reversal across years, while estimating expected genotypic sex ratios for ungenotyped years by leveraging data from years with complete genetic information. We investigated the environmental drivers of these probabilities by incorporating both maximum and minimum daily temperatures. While maximum temperatures have been historically used as an index for thermal condition in this species [35–38,41], we expanded our models to include minimum temperatures, which usually occur at night and represent a critical, unbuffered thermal period for reptile physiology [51–54].

If temperature-dependent phenotypic sex ratio variation is driven by annual genotypic sex ratio variation, genotypic sex ratios are likely themselves temperature-dependent. We modelled the probability of being a genetic male (*q*), when born from mother *m*, during year *y* at population *s,* as:

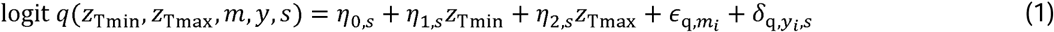

where *z_Tmin_* and *z_Tmax_* are the z-transformed mean minimum and maximum temperatures during the sex-determining period. The z-transformations were performed at the population level, accounting for biological and habitat differences that likely led to variation in how air temperature aligned with experienced microclimates [41]. The *η*-terms were parameters to be estimated, describing how temperature affected the genotypic sex ratios at each site. We included normally-distributed random effects for maternal differences (*ɛ_q,mi_*) and unexplained year nested within population variation (*δ_q,yi,s_*).

Similarly, the probability of an offspring of genotypic sex *g* experiencing sex reversal (*r_g_*) was modelled as:

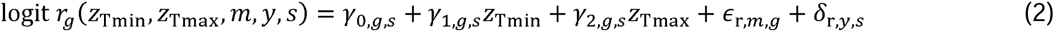

where *γ*-parameters describe temperature effects on sex reversal likelihood for each genetic sex and population. *ɛ_r,m,g_* and *δ_r,y,s_* are normally-distributed random effects describing maternal effect and population-specific annual variation in sex reversal, respectively.

To ensure robust estimates of *q* and *r_g_*, our models incorporated laboratory-derived misclassification rates from Hill et al. [39], accounting for uncertainty introduced by the phenotypic sex assessment process itself. We defined *e_M_* and *e_F_* as the probabilities of incorrectly assigning the male and female phenotypes, respectively (likelihood equations in Table S1).

Our primary goal was to estimate reversal rates for both genetic sexes: *r*_XX_ for genetic females (XX) developing as phenotypic males (M); and *r*_XY_ for genetic males (XY) developing as phenotypic females (F). Table 1 provides the probabilities for the four genotype-phenotype combinations. We applied these probabilities differently depending on the data available for each year. For years where we had complete information for each mother on the number of offspring observed in each genotype-phenotype category (denoted by *n_g,p,m,y_*), the log-likelihood of each combination given our model was:

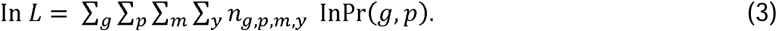

**Table 1:** Probabilities for the four possible genotype-phenotype combinations observed in offspring. Probabilities are calculated using the likelihood of a genotype (*q*), sex reversal given genotype (*rg*), and phenotypic misclassification (*e_M_, e_F_*). Note that *q* and *r_g_* vary between mothers, sites and years, and depend on temperature (see Eqns. 1 and 2).

| Genotype, $g$ | Phenotype, $p$ | Probability offspring has genotype/phenotype, $\Pr(g, p)$ |
| --- | --- | --- |
| XY | M | $q(r_{XY}e_F + (1 - r_{XY})(1 - e_M))$ |
| XY | F | $q(r_{XY}(1 - e_F) + (1 - r_{XY})e_M)$ |
| XX | M | $(1 - q)(r_{XX}(1 - e_M) + (1 - r_{XX})e_F)$ |
| XX | F | $(1 - q)(r_{XX}e_M + (1 - r_{XX})(1 - e_F))$ |

Conversely, for years where there was only information about phenotypic counts per mother but not genetic sex information (*n_p,m,y_*), the model inferred sex reversal by assuming genotypic sex ratios exhibited random variation consistent with genotyped years (defined by *δ_q,yi,s_* in Eq. 1). In this case, the probability an offspring exhibited phenotype *p*, was the probability it was a genetic male and exhibited phenotype *p* plus the probability it was a genetic female and exhibited phenotype *p*, which gives:

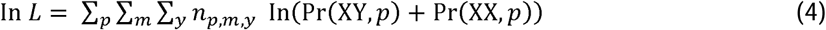

Bayesian estimates of model parameters were performed in R (v4.3.2) [55], using Stan via the *rstan* package (v2.32.3) [56]. The model was fit simultaneously to the phenotypic, genotypic, and misclassification data. Probabilities for each of the four genotype-phenotype pairings (Table 1) were incorporated into the likelihood function for genotyped individuals (Eq. 3) and non-genotyped individuals (Eq. 4). Misclassification rates were extracted from Table S2. Wide prior distributions were used to ensure parameter estimates were data-driven, and inference was based on 2,000 posterior samples following a 1,000-sample burn-in, after verifying chain convergence. Full model implementation details, including the annotated R and Stan scripts used for this analysis, are provided in the Supplementary Materials.

### Modelling temperature effect on phenotypic sex ratios

Having established the thermal dependencies of genotypic sex and sex reversal, we tested whether these explained the long-term phenotypic skews observed in wild populations. To achieve this, we updated previous phenotypic analyses [37] by expanding the dataset to 23 years (N = 3,369 offspring), ensuring we had phenotypic confirmation of sex ratio skews for the recent cohorts targeted for genetic sexing.

We modelled the number of phenotypic male offspring per clutch using a binomial generalized linear mixed model (GLMM) with a logit link function. Fixed effects included site, z-transformed values of mean maximum and minimum temperatures, and their interactions. Year nested within population was included as a random effect to account for inter-annual variability; maternal identity was excluded due to negligible variance. To directly test if sex reversal was responsible for phenotypic biases, we considered including annual sex reversal frequencies as a model covariate. However, empirical genetic data were only available for a small subset of years — limiting statistical power for a direct test — and the wide credible intervals associated with Bayesian estimates for the remaining ungenotyped cohorts rendered them unsuitable as firm predictors.

Parameter uncertainty was quantified using Bayesian methods, simulating 10,000 posterior draws assuming uninformative priors. Inference was based on the posterior probability of a positive regression coefficient: posterior values near 0 or 1 indicated strong evidence of a negative and positive directional effect, respectively. The analysis was performed in R, with the statistical model built using the package *lme4* [57] and posterior parameters simulated using the *arm* package [58].

### Identification of sex reversed adult males

To assess if XX males — the only sex-reversal type previously demonstrated in laboratory tests [39] — persist into adulthood, we sampled most adult phenotypic males (∼90–95%) in 2022 from the lowland (N = 59) and highland (N = 144) populations. Captured individuals were tail-clipped as a source of DNA, toe-clipped for identification (if not done previously as offspring), and immediately released. Phenotypic sex of adult males was assessed by hemipenis eversion, a method considered unambiguous in sexually mature individuals. Genotypic sex was assessed using the same PCR protocol as in offspring samples.

To increase our survey effort and geographic coverage, we also sampled phenotypic males from 6 additional populations (Figure S1). This includes 20 individuals from a low-altitude population (Bicheno: 41°52’ S, 148°18’ E, 4 m a.s.l.); 60 individuals from three mid-altitude populations (20 from Lagoon of Islands: 42°06’ S, 146°55’ E, 758 m a.s.l.; 20 from Tooms Lake: 42°13’ S, 147°46’ E, 460 m a.s.l.; and 20 from Lake Echo: 42°13’ S, 146°38’ E, 850 m a.s.l.); and 38 individuals from two other high-altitude populations (18 from kunanyi/Mt Wellington: 42°53’ S, 147°14’ E, 1,000 m a.s.l.; and 20 from Ben Lomond: 41°30’ S, 147°39’ E, 1,160 m a.s.l.).

## 3. Results

### Phenotypic and genotypic data, sex reversal and misclassification rates

Across 23 years (2000 to 2022), the phenotypic sex of 11,486 *C. ocellatus* offspring from 1,579 females was described at two long-term study sites (Table S2). Between 46 and 123 pregnant females from the highland population were captured annually, producing an average clutch of 3.62 (±0.077) offspring. At the lowland population, between 48 and 111 pregnant females were captured annually, producing an average clutch of 2.37 (±0.024) offspring. Over this period, a weak phenotypic male-bias was evident in both populations, slightly more pronounced in the lowland (Figure 2a). The lowland population had greater annual variation in phenotypic sex ratios, with several years showing a strong male skew (>58% males e.g., 2001, 2003, 2004, 2013, 2019) the highest being 2003 with >65% males, but also a few years showing female-skews (e.g., 2000, 2005, 2018; 45-46% males).

We genetically sexed 2,582 offspring (highland: 1,516; lowland: 1,066) across the study period (Figure 2b; Table S2). Based on laboratory misclassification rates [39] and our 23-year phenotypic dataset, our model estimated that phenotypic sex misclassification is low: a 0.2% [95% CI: 0.0,0.7%] probability of falsely scoring a phenotypic male as a female (e_M_) and a 2.4% [95% CI: 1.3,3.8%] probability for the converse (e_F_). Frequencies of XX males were generally low in both populations (mean highland: 1.2%; mean lowland: 4.1%; Figure 2c, Table S2). However, they reached notable peaks in 2019 and 2022 where they represented 23.5% and 11% of all XX individuals born in the lowland population respectively (Figure 2c), substantially exceeding the 2.4% misclassification rate for males. Despite 35 XX males being born in the period from 2018-2021 in the lowland population (Table S2), persistence to adulthood appears limited. Specifically, among 321 wild-caught adult males sampled across eight populations in 2022, only three were identified as sex-reversed (one from the long-term lowland population and two from Tooms Lake, a mid-altitude site). Although generally rare, XY females were consistently identified among newborns (mean highland: 2.1%; mean lowland: 0.9%; Figure 2c, Table S2), exceeding the 0.2% misassignment rate for phenotypic females in most years.

### Modelling genotype, sex reversal and temperature dependence

We found no evidence that temperature influenced the underlying genotypic sex ratio at either site (Table 2; Figure 3a). However, the probability of sex reversal in genetic females (producing XX males) varied along a reaction norm, with significantly higher rates at lower temperatures (Table 2). This effect was stronger in the lowland population, where both minimum and maximum temperatures contributed to the fit. By integrating these thermal dependencies, the model successfully reconstructed the observed peaks of XX males in 2019 and 2022 (Figure 3b). Furthermore, the model attributes the strong phenotypic male bias observed in 2003 and 2004 to sex reversal, providing a mechanistic explanation for these years’ phenotypic bias (Figure 2a, 3b). In contrast, reversal in genetic males (XY females) showed no association with temperature, resulting in consistently low, baseline reversal rates across the study period (Table 2; Figure 3c).

**Figure 3:**
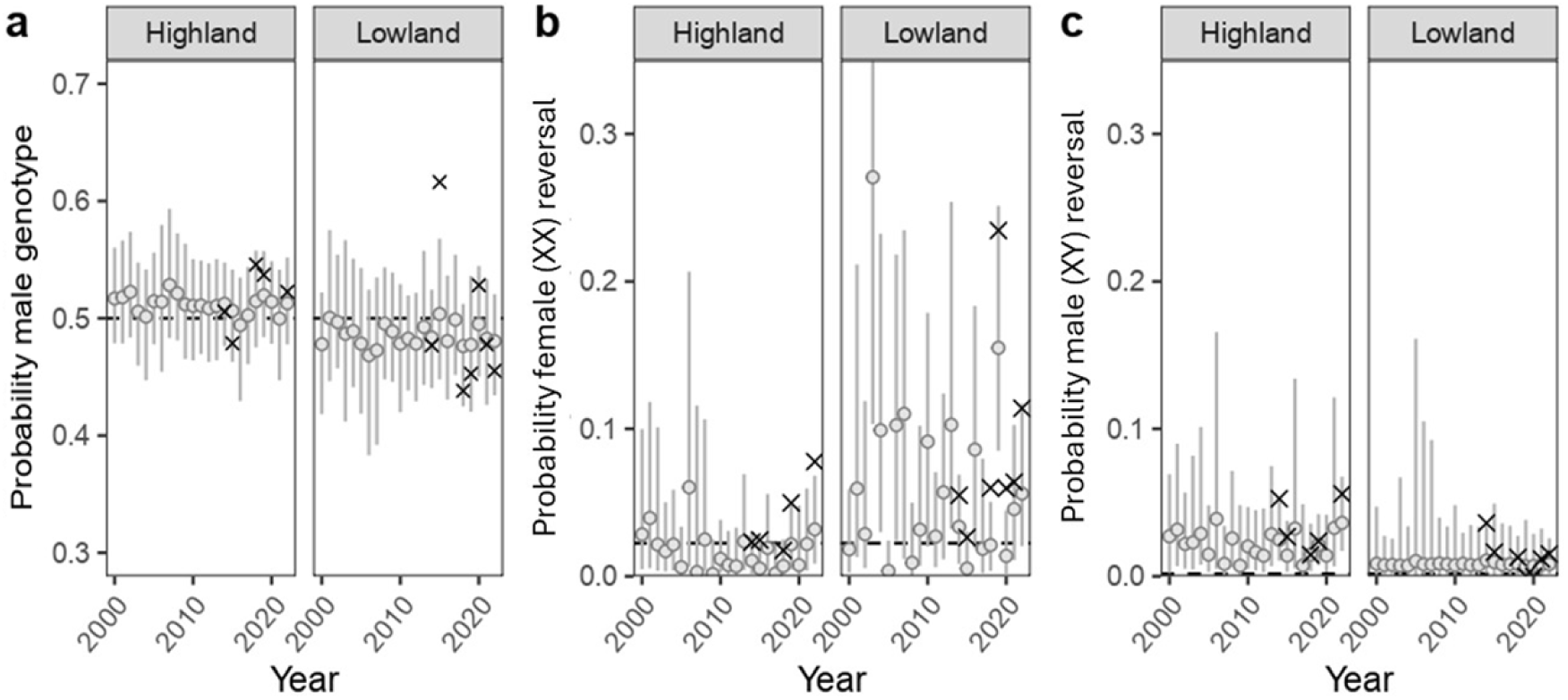
Observed and predicted genotypic sex ratios and sex reversal rates. a) Yearly estimates for the proportion of genetic males. b) Genetic female reversal (XX males; *r_XX_*). c) Genetic male reversal (XY males; *r_XY_*). Circles and error bars represent yearly estimates and 95% credible intervals (CIs), while crosses denote observed proportions. Dashed lines in (b) and (c) mark the probability of phenotypic misclassification (*e_F_* and *e_M_*, respectively), which positively biases the observed crosses relative to the estimates. *Note: The 2015 lowland male skew in (a) is a sampling artifact caused by disproportionate pre-screening loss of phenotypic females*.

**Table 2:** Parameter estimates and credibility associated with temperature dependencies for genotypic sex and sex reversal. Variables include the probability offspring being genotypically male 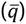, and the probability of sex reversal in genetic males 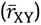 and females 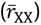. Estimates represent posterior medians [95% credible intervals] calculated separately for highland (HE) and lowland (LE) populations at site-specific mean temperatures. The table also provides the credibility of positive associations with minimum and maximum temperatures, alongside brief conclusions.

| Variable | Estimate [95%CI] | Condition | Credibility | Conclusion |
| --- | --- | --- | --- | --- |
| **Genotypic sex ratio (probability offspring are male)** |  |  |  |  |
| $\bar{q}(\text{HE})$ | 0.512 [0.495,0.530] | | | unbiased sex ratio at highland |
| | | $\eta_{1,\text{HE}} > 0$ | 0.432 | no minimum temperature effect |
| | | $\eta_{2,\text{HE}} > 0$ | 0.539 | no maximum temperature effect |
| $\bar{q}(\text{LE})$ | 0.486 [0.458,0.508] | | | unbiased sex ratio at lowland |
| | | $\eta_{1,\text{LE}} > 0$ | 0.606 | no minimum temperature effect |
| | | $\eta_{2,\text{LE}} > 0$ | 0.616 | no maximum temperature effect |
| **Phenotypic sex reversal (probability offspring phenotype differs from genotype)** |  |  |  |  |
| $\bar{r}_{XY}(\text{HE})$ | 0.021 [0.010,0.038] | | | low reversal of males at highland |
| | | $\gamma_{1,XY,\text{HE}} > 0$ | 0.331 | no minimum temperature effect |
| | | $\gamma_{2,XY,\text{HE}} > 0$ | 0.149 | no maximum temperature effect |
| $\bar{r}_{XY}(\text{LE})$ | 0.009 [0.003,0.020] | | | very low reversal of males at lowland |
| | | $\gamma_{1,XY,\text{LE}} > 0$ | 0.549 | no minimum temperature effect |
| | | $\gamma_{2,XY,\text{LE}} > 0$ | 0.510 | no maximum temperature effect |
| $\bar{r}_{XX}(\text{HE})$ | 0.012 [0.004,0.026] | | | very low reversal of females at highland |
| | | $\gamma_{1,XX,\text{HE}} > 0$ | 0.156 | no minimum temperature effect |
| | | $\gamma_{2,XX,\text{HE}} > 0$ | 0.097 | negative maximum temperature effect |
| $\bar{r}_{XX}(\text{LE})$ | 0.041 [0.012,0.074] | | | low reversal of females at low elevation |
| | | $\gamma_{1,XX,\text{LE}} > 0$ | 0.003 | negative minimum temperature effect |
| | | $\gamma_{2,XX,\text{LE}} > 0$ | 0.093 | negative maximum temperature effect |

### Modelling temperature dependence of phenotypic sex ratios

Having established via our Bayesian model that genotypic sex ratios are broadly at parity with no significant environmental association (Table 2, Figure 3a), we next examined whether fluctuations in phenotypic sex ratios were congruent with temperature-induced sex reversal. Confirming this mechanism requires demonstrating that long-term phenotypic skews align closely with the same fine-scale thermal cues known to drive reversal.

Consistent with the sex-reversal mechanism, our model reveals that years with colder minimum temperatures were significantly associated with an increased proportion of phenotypic males at the lowland site (credibility positive effect = 0.002; Figure S2). In contrast, there was little effect of minimum temperatures on phenotypic sex ratios at the highland site (credibility positive effect = 0.403; Figure S2). Maximum temperature had a negligible effect on phenotypic sex ratios at both sites (lowland: credibility positive effect = 0.430; highland: credibility positive effect = 0.539; Figure S2).

## 4. Discussion

Our study is the first demonstration of sex reversal in wild viviparous reptiles, reinforcing previous observations under experimental laboratory conditions [39,59]. By establishing the occurrence of this phenomenon under natural thermal regimes, our findings confirm that sex reversal is not merely a laboratory artifact and offer a critical foundation to explore its broader ecological and evolutionary implications. Our study contributes to the growing body of evidence of sex reversal across vertebrates [10,25,26], and particularly in reptiles [24,29,60], proving that even viviparous species with GSD are susceptible to thermal influence. Furthermore, we showed that cold conditions increased female-to-male sex reversal (XX males), offering a potential mechanism for the male-biased phenotypic sex ratio observed during cold years at the lowland population [36–38]. Because our analysis confirmed that the underlying genotypic sex ratios remain stable across these thermal extremes, we can confidently infer that environmentally induced sex reversal is a primary driver of these population-level phenotypic skews, matching theoretical predictions [7,9].

Previous studies have linked *C. ocellatus* sex ratios and reversal to lower maximum daytime temperatures at the lowland site [35–37,39]. Yet, our model suggests that low daily minimum temperatures, usually occurring at night, may act as an additional, critical predictor. This dependence on both thermal extremes is consistent with a X-dosage threshold model for sex determination (Figure 1) [16,17]. Under this model, cold conditions promote sex reversal by inhibiting the accumulation of sex-determining factors required for female phenotypic development. Consequently, the probability of sex reversal depends on the cumulative time embryos spend at sub-threshold temperatures, a duration that is synergistically affected by both temperature extremes. Minimum temperatures influence the expression of embryonic sex-determining factors during nocturnal retreat, a period when females have little to no control over their body temperature. Conversely, maximum temperatures represent maternal opportunity to behaviourally thermoregulate, which interplays with ambient daytime conditions to dictate embryo’s thermal experience. Collectively, these results underscore the physiological importance of nighttime thermal conditions in reptiles [51–54]. Interestingly, the historical association between maximum temperatures and phenotypic sex ratios appears to have weakened over the course of our extended dataset. This is potentially due to forest encroachment at the lowland site, which could decouple measured macroclimates from the microclimates experienced by females. Ultimately, sex reversal in *C. ocellatus* seems to parallel the TSD pattern typical of turtles, where cold favours male development [61,62], and aligns with observations of cold-induced sex reversal in *Acritoscincus duperreyi* [63].

Population differences in the frequency of XX males present an apparent ecological paradox: if low temperatures drive sex reversal, why does the much colder highland environment produce fewer XX males? The resolution probably lies in the divergent selective pressures on each population. In the lowlands, cold years disproportionately reduce female fitness (e.g., delayed maturation and at a reduced size), favouring temperature-sensitive sex determination [3,38]. Conversely, the high thermal variability and lack of sex-specific fitness benefits in the highlands select against this thermal sensitivity to maintain balanced sex ratios [37,38,41]. This reduced sensitivity likely involves both genetic adaptation and compensatory maternal behaviour. Although cold conditions induce reversal in both populations with similar laboratory reaction norms, there was a non-significant trend towards lower frequencies of XX reversal in the highland population [39]. Coupled with the low frequencies of XX reversals observed in the wild highland population, this points to an underlying genetic divergence in the mode of sex determination. Assuming a X-dosage mode of sex determination as depicted in Figure 1, this divergence may include shifts in (i) the threshold for female phenotypic development, (ii) the baseline production of sex-determining factor, (iii) the reaction norms for factor expression, and/or (iv) the temperature of peak expression. Behaviourally, highland females compound this divergence by exhibiting a more intense basking behaviour [40,47] and potentially optimising their retreat selection, actively reducing embryonic exposure to cold conditions. Yet, the paradox does not end with these potential adaptations; it is further compounded by a striking contradiction between laboratory and field data. Specifically, sex reversal occurs more frequently under controlled laboratory conditions than in wild highland individuals, even though laboratory temperatures were generally warmer than their natural highland habitat (Figure S3) [39]. This discrepancy highlights the broader challenge of replicating natural environmental and thermal dynamics in controlled setups [64] — especially in viviparous reptiles — while also providing a unique opportunity to experimentally isolate the specific mechanisms driving sex reversal in this species.

The unexpected occurrence of XY females in *C. ocellatus*, while rare and requiring further experimental confirmation, raises the possibility that bidirectional sex reversal occurs in reptiles — a phenomenon previously documented only in amphibians and teleost fish [20,65,66]. In the lowland population, XY females would offer a novel mechanism to explain phenotypically female-biased years [35–38], specifically when XY reversal rates outpaces XX reversals. Previously, these phenotypic biases were attributed to consistently female-bias genetic cohorts, caused either by the overproduction of genetic females at conception or by XX males exclusively siring XX offspring [23,39]. However, the broadly balanced genetic sex ratios observed here suggest that lowland female skews could be driven by XY reversal, alongside contributions from minor interannual genetic fluctuations. In the highland population, phenotypic sex parity could, in part, be maintained, by XY females balancing the production of XX males. Under the threshold model for sex determination, the observed increased XY female and suppressed XX male prevalence would match with a lowered threshold for female development in the highland population (as in Figure 1a). We found no direct thermal link to XY reversal, reflecting either limited statistical power or an alternative, potentially non-thermal mechanisms driving it (e.g., stochastic developmental noise intrinsic to the threshold model) [67]. Bidirectional sex reversal opens compelling new avenues for investigating the molecular and developmental pathways of sex determination.

A pressing question following the discovery of sex reversal in *C. ocellatus* offspring concerns whether these individuals persist into the adult breeding population, where skewed sex ratios can negatively impact population dynamics and genetic diversity [6–9]. Our results indicate that adult XX males are rare across all sampled populations, even in the lowland population, where high frequencies of reversals were detected in a recent offspring cohort. This limited recruitment contrasts sharply with other reptiles, such as *Pogona vitticeps,* where 6–27% of the adult females are sex reversed [31], and *A. duperreyi*, which sees up to 18% adult XX males in some populations [30]. This absence of adult XX males could be explained by phenotypic developmental delays. For instance, delays in hemipenis regression, as observed in species like *Barisia imbricata* [68], could result in transient phenotypic mismatches at birth that later resolve in XX females. Alternatively, sex reversal in *C. ocellatus* may simply not confer a fitness advantage and instead carry significant costs, potentially mirroring the physiological stress and reduced survival observed in sex-reversed juvenile *Rana dalmatina* [18,19]. Resolving whether this absence stems from transient developmental artifacts or genuine fitness costs remains a priority for future research.

However, if true sex reversals do indeed suffer high mortality, their continued presence in the wild raises intriguing evolutionary questions. In such a scenario, the rarity of these reversal events may mean that selection against the underlying genetic architecture is too weak to eliminate it, effectively behaving as neutral, allowing the trait to persist through the sheltering of deleterious alleles in heterozygotes [69], or via spatially and temporally variable selection [70].

Ultimately, whether sex reversal will be maintained in *C. ocellatus* likely depends on the progression of anthropogenic climate change. Under projected baseline warming, the frequency of cold reversal-inducing conditions is expected to decline [71], potentially diminishing the occurrence of XX reversal over time. However, climate change is equally characterized by increased thermal instability, which could promote localized “cold snaps” during critical embryonic windows [72]. These extreme events could still trigger sudden, maladaptive sex ratio skews, threatening population viability in an otherwise warming world [73]. Consequently, evaluating the interplay between fitness costs of sex reversal and this shifting thermal landscape will be essential for assessing the resilience of *C. ocellatus* in an increasingly unpredictable world.

