## Supplementary Material for "Thermally driven sex reversal reveals divergent sex determination dynamics in wild viviparous reptile populations"

SUPLEMENTARY MATERIAL

**Table S1:** Probabilities of phenotypic assessment outcomes conditional on genotype. Data are derived from Hill et al. [39] using a blind repeat-assessment protocol with a third 'tie-breaker' trial by assessor EW. robabilities are calculated using the likelihood of a male genotype ($q$), sex reversal given genotype ($r_{g}$), and phenotypic misclassification ($e$), which differs depending on the true phenotypic sex of the offspring (M,F). Note that the $q$ and $r$ vary between mothers, sites and years, and depend on temperature (see Eqns. 2 and 3).

| Genotype | Phenotypic assessment | Probability |
| --- | --- | --- |
| XY | 2M (Consistent male) | $q(r_{\mathrm{XY}}e_{F}+\left( 1-r_{\mathrm{XY}} \right)\left( 1-e_{M} \right))$ |
| XY | 2F (Consistent female) | $q(r_{\mathrm{XY}}{(1-e}_{F})+\left( 1-r_{\mathrm{XY}} \right)e_{M}$) |
| XX | 2M (Consistent male) | $(1-q)(r_{\mathrm{XX}}(1-e_{M})+\left( 1-r_{\mathrm{XX}} \right)e_{F})$ |
| XX | 1M,2F (Discrepancy resolved as female) | $(1-q)(r_{\mathrm{XX}}e_{M}+\left( 1-r_{\mathrm{XX}} \right)\left( 1-e_{F} \right))$ |

**Table S2:** Annual breakdown of gravid females captured and offspring sex distributions (phenotypic vs. genetic) by population. Counts are provided for phenotypic males and females, as well as for specific genotype-phenotype combinations (including sex-reversed XX males and XY females) during years of genetic assessment. Note: Genotyping coverage was high across most years tested (>95%), with a minimum coverage of 78% in the 2015 lowland cohort.

| Population | Year | Mothers | Phenotypic males | Phenotypic females | XY males | XX females | XX males | XY females |
| --- | --- | --- | --- | --- | --- | --- | --- | --- |
| Highland | 2000 | 90 | 204 | 173 | - | - | - | - |
| Highland | 2001 | 115 | 240 | 198 | - | - | - | - |
| Highland | 2002 | 83 | 166 | 129 | - | - | - | - |
| Highland | 2003 | 102 | 173 | 178 | - | - | - | - |
| Highland | 2004 | 100 | 151 | 165 | - | - | - | - |
| Highland | 2005 | 95 | 169 | 151 | - | - | - | - |
| Highland | 2006 | 87 | 141 | 122 | - | - | - | - |
| Highland | 2007 | 83 | 154 | 111 | - | - | - | - |
| Highland | 2008 | 67 | 123 | 96 | - | - | - | - |
| Highland | 2009 | 48 | 96 | 88 | - | - | - | - |
| Highland | 2010 | 46 | 82 | 79 | - | - | - | - |
| Highland | 2011 | 49 | 91 | 85 | - | - | - | - |
| Highland | 2012 | 51 | 91 | 92 | - | - | - | - |
| Highland | 2013 | 70 | 140 | 134 | - | - | - | - |
| Highland | 2014 | 71 | 134 | 136 | 126 | 127 | 3 | 7 |
| Highland | 2015 | 62 | 113 | 122 | 109 | 119 | 3 | 3 |
| Highland | 2016 | 100 | 174 | 205 | - | - | - | - |
| Highland | 2017 | 115 | 229 | 237 | - | - | - | - |
| Highland | 2018 | 93 | 169 | 153 | 135 | 112 | 2 | 2 |
| Highland | 2019 | 123 | 244 | 209 | 205 | 172 | 9 | 5 |
| Highland | 2020 | 118 | 205 | 184 | - | - | - | - |
| Highland | 2021 | 110 | 175 | 192 | - | - | - | - |
| Highland | 2022 | 119 | 200 | 178 | 186 | 166 | 14 | 11 |
| Lowland | 2000 | 92 | 89 | 106 | - | - | - | - |
| Lowland | 2001 | 101 | 135 | 92 | - | - | - | - |
| Lowland | 2002 | 90 | 115 | 90 | - | - | - | - |
| Lowland | 2003 | 105 | 141 | 71 | - | - | - | - |
| Lowland | 2004 | 93 | 128 | 90 | - | - | - | - |
| Lowland | 2005 | 111 | 126 | 156 | - | - | - | - |
| Lowland | 2006 | 93 | 109 | 113 | - | - | - | - |
| Lowland | 2007 | 80 | 89 | 87 | - | - | - | - |
| Lowland | 2008 | 109 | 128 | 120 | - | - | - | - |
| Lowland | 2009 | 99 | 128 | 116 | - | - | - | - |
| Lowland | 2010 | 106 | 137 | 121 | - | - | - | - |
| Lowland | 2011 | 94 | 114 | 118 | - | - | - | - |
| Lowland | 2012 | 87 | 96 | 96 | - | - | - | - |
| Lowland | 2013 | 81 | 120 | 80 | - | - | - | - |
| Lowland | 2014 | 73 | 87 | 92 | 80 | 86 | 5 | 3 |
| Lowland | 2015 | 50 | 68 | 56 | 60 | 37 | 1 | 1 |
| Lowland | 2016 | 48 | 64 | 54 | - | - | - | - |
| Lowland | 2017 | 61 | 85 | 67 | - | - | - | - |
| Lowland | 2018 | 87 | 97 | 114 | 77 | 94 | 6 | 1 |
| Lowland | 2019 | 71 | 90 | 63 | 67 | 62 | 19 | 0 |
| Lowland | 2020 | 62 | 81 | 69 | 75 | 63 | 4 | 0 |
| Lowland | 2021 | 76 | 92 | 90 | 85 | 88 | 6 | 1 |
| Lowland | 2022 | 56 | 75 | 71 | 65 | 70 | 9 | 1 |

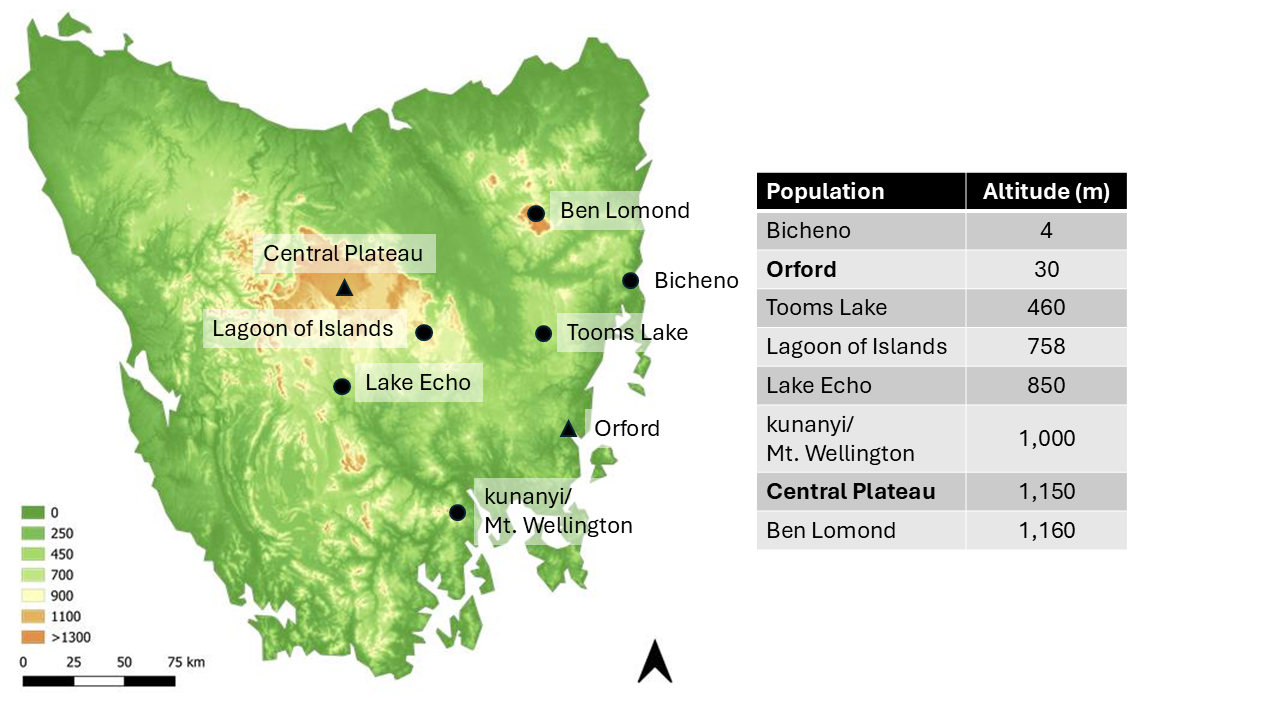

**Figure S1:** Geographical distribution and elevation of *C. ocellatus* populations assessed for adult sex reversal. Symbols on the map correspond to sampling locations: triangles denote the two long-term monitoring sites (lowland = Orford and highland = Central Plateau), while circles represent additional survey populations. The accompanying table lists the altitude (m.a.s.l.) for each site.

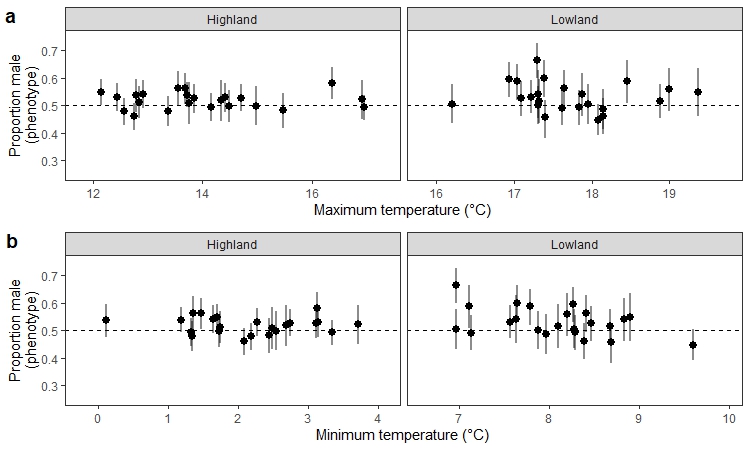

**Figure S2:** Average phenotypic male proportion per clutch in each population and year considering a) mean maximum temperature and b) mean minimum temperature during the critical sex-determining period of the respective year. Points describe the average proportion of males per clutch at each year’s temperature and bars represent 95% confidence intervals. Dashed line describes the expected 50:50 proportion.

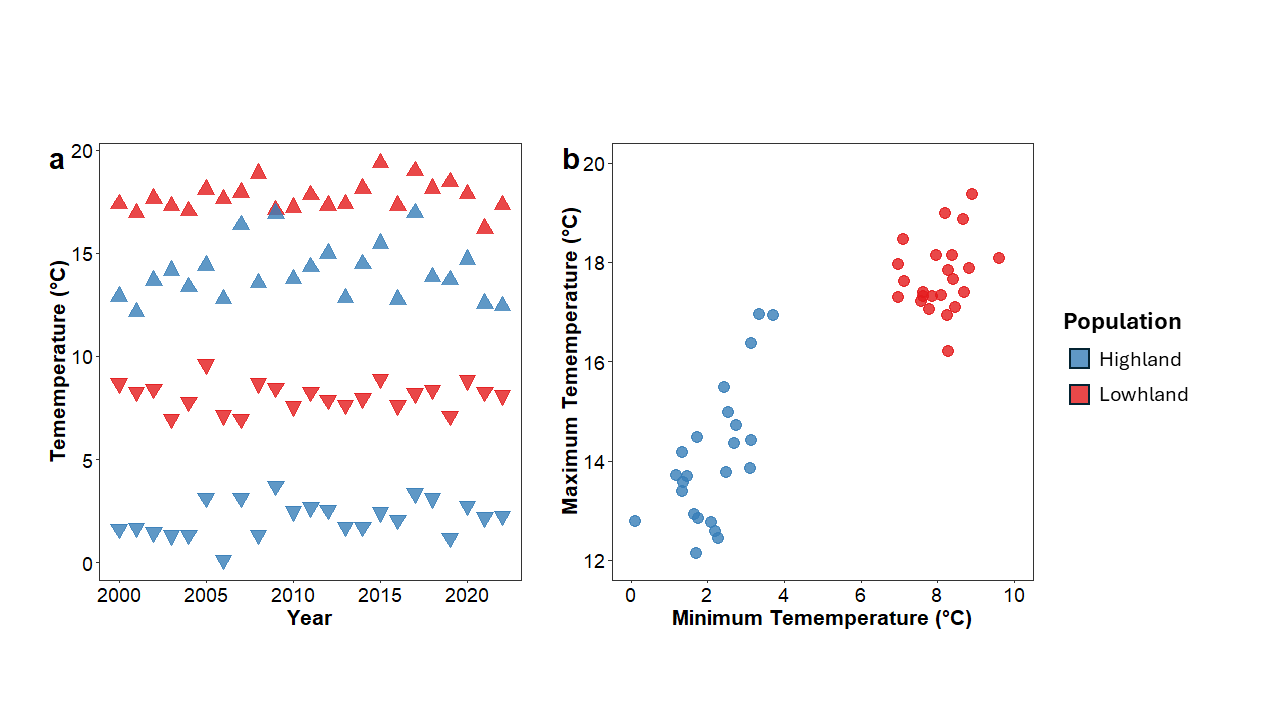

**Figure S3:** Distinct thermal environments of the highland and lowland *C. ocellatus* populations. (a) Annual variation in environmental temperatures from 2000 to 2022. Upward-pointing triangles denote maximum temperatures, while downward-pointing triangles denote minimum temperatures for the highland (blue) and lowland (red) populations. (b) Scatter plot illustrating the relationship between maximum and minimum temperatures across the same period.
